# Data-driven clustering reveals a link between symptoms and functional brain connectivity in depression

**DOI:** 10.1101/267591

**Authors:** Luigi A. Maglanoc, Nils Inge Landrø, Rune Jonassen, Tobias Kaufmann, Aldo Cordova-Palomera, Eva Hilland, Lars T. Westlye

**Affiliations:** Clinical Neuroscience Research Group, Department of Psychology, University of Oslo, Oslo, Norway; NORMENT, KG Jebsen Centre for Psychosis Research, Division of Mental Health and Addiction, Oslo University Hospital & Institute of Clinical Medicine, University of Oslo, Norway; Division of Psychiatry, Diakonhjemmet Hospital, Oslo, Norway; Department of Pediatrics, Stanford University School of Medicine, Stanford, CA; Department of Psychology, University of Oslo, Oslo, Norway

**Keywords:** depression, symptom-based clustering, fMRI, heterogeneity, functional connectivity, anxiety

## Abstract

**Background:** Depression is a complex disorder with large inter-individual variability in symptom profiles that often occur alongside symptoms of other psychiatric domains such as anxiety. A dimensional and symptom-based approach may help refine the characterization and classification of depressive and anxiety disorders and thus aid in establishing robust biomarkers. We assess the brain functional connectivity correlates of a symptom-based clustering of individuals using functional brain imaging data.

**Methods:** We assessed symptoms of depression and anxiety using Beck’s Depression and Beck’s Anxiety inventories in individuals with or without a history of depression, and high dimensional data clustering to form subgroups based on symptom profiles. To assess the biological relevance of this subtyping, we compared functional magnetic resonance imaging-based dynamic and static functional connectivity between subgroups in a subset of the total sample.

**Results:** We identified five subgroups with distinct symptom profiles, cutting across diagnostic boundaries and differing in terms of total severity, symptom patterns and centrality. For instance, *inability to relax, fear of the worst, and feelings of guilt* were among the most severe symptoms in subgroup 1, 2 and 3, respectively. These subgroups showed evidence of differential static brain connectivity patterns, in particular comprising a fronto-temporal network. In contrast, we found no significant associations with clinical sum scores, dynamic functional connectivity or global connectivity measures.

**Conclusion:** Adding to the ongoing pursuit of individual-based treatment, the results show subtyping based on a dimensional conceptualization and unique constellations of anxiety and depression symptoms is supported by distinct brain static functional connectivity patterns.

## Introduction

Major depressive disorder is among the leading contributors to years lived with disability,^1^ and the leading cause in 56 countries.^2,3^ Although several brain regions have been implicated in the pathophysiology of depression including the subgenual anterior cingulate,^4^ global efforts for identifying sensitive, specific and clinically predictive brain correlates of mood disorders have still not succeeded.^5,6^ One reason for the lack of robust imaging-based characteristics is that depression is a heterogeneous construct with regards to symptom constellation. For example, based on 12 items from the Quick Inventory of Depressive Symptomatology (QIDS-16), Fried and Nesse^7^ found evidence of 1030 unique symptom profiles among 3703 depressed outpatients. Remarkably, the most common symptom profile had a frequency of < 2%, and > 80% were shared by five or fewer subjects and almost 50% by only one individual. Furthermore, depression and anxiety symptoms often co-occur, exemplified by 75% of individuals with a depressive disorder in the Netherlands Study of Depression and Anxiety (NESDA) study also fulfilling the criteria for an anxiety disorder.^8^ Adding to the complexity, individuals of the general, healthy population from time to time also experience depression symptoms at various degrees.

Methodological variability is another reason for the mixed brain imaging findings in depression, especially for functional MRI-based measures of brain activation^6^ and connectivity.^9^ The functions of a healthy mind are supported by the continuous cross talk between different brain regions.^10^ Dysregulations in this fine-tuned and complex interplay may increase susceptibility for mental disorders^11^. Supporting the conceptualization of depression as a network-based disorder, fMRI-based functional connectivity (FC) studies have implicated large-scale brain network dysfunction in depression.^9^ Whereas previous studies have primarily reported results from various estimates of static FC (sFC; the temporal correlation between two brain regions across the entire time-series), there is an increasing awareness of the relevance of dynamic FC (dFC; the variability in the temporal correlations across the time-series).^12,13^ Interestingly, sFC and dFC capture distinct properties of brain network dynamics,^14,15^ and may therefore provide complementary information in depression.^16^

Here, in order to address symptom heterogeneity in depression, we used high dimensional data-driven clustering (HDDC)^17^ based on item scores on the Beck’s Depression (BDI-II) and Beck’s Anxiety (BAI) inventories to identify groups of individuals with distinct symptom profiles among 1084 subjects with or without a history of a diagnosis of depression. In order to assess the brain system-level relevance of the symptom-based subgroups, we compared measures of fMRI-based static and dynamic connectivity between groups in a subset of 251 individuals using network-based statistics.

## Methods and Materials

### Sample

In the total sample (N = 1084), 605 individuals with a history of major depressive episodes (MDE) and individuals with no history of an MDE (N = 437) were included (Table 1), drawn from four research projects at the Clinical Neuroscience Research Group, Department of Psychology, University of Oslo (see the Supplemental Methods). An MRI-subsample of 251 participants (Table 2) was drawn from one of these research projects (see the Supplemental Methods). Individuals with a history of depression were diagnosed using the Structural Interview for DSM-IV (SCID-I)^18^ in one of the sub-studies and Mini International Neuropsychiatric Interview (M.I.N.I 6.0)^19^ in the other three, and were mainly recruited from outpatient clinics. Individuals with no history of depression were recruited by posters, advertisements in the local newspaper, and social media. The presence of other major Axis I psychiatric disorders and number of lifetime MDEs were assessed for all participants based on the M.I.N.I 6.0 (or SCID-I in one of the sub studies). Current selective serotonin reuptake inhibitor (SSRI) use was evaluated through a semi-structured interview. Individuals with a history of neurological disorders and MRI contraindications (for the MRI-sample) were excluded. The study was approved by the Regional Ethical Committee of South-Eastern Norway (REK Sør-Øst) and all participants provided informed consent prior to enrolment.

**Table 1.** Demographics for the total sample

|  | Individuals with<br>no history of<br>depression<br>(n = 437) |  | Individuals with a<br>history of<br>depression<br>(n = 605) |  | *p* |
| --- | --- | --- | --- | --- | --- |
|  | M | SD | M | SD |  |
| Gender (female) | n = 287 |  | n = 468 |  | <.001 |
| Age | 33.90 | 13.35 | 39.45 | 12.92 |  |
| Depression symptoms |  |  |  |  | <.001 |
| BDI-II | 4.45 | 5.50 | 14.09 | 11.22 |  |
| Anxiety symptoms |  |  |  |  | <.001 |
| BAI | 3.15 | 4.11 | 8.68 | 8.36 |  |
| Other |  |  |  |  | <.001 |
| History of anxiety disorder | n = 7 |  | n = 153 |  | <.001 |
| History of (hypo)mania | n = 0 |  | n = 63 |  | <.001 |
| No. of depressive episodes | 0 |  | 4.12 | 6.49 | <.001 |
| Currently medicated (SSRI) | n = 0 |  | n = 164 |  | <.001 |
| Currently unmedicated (SSRI) | n = 0 |  | n = 383 |  | <.001 |

An additional 42 cases were left out from the table because it could not be determined from the records to what extent they have had a depressive episode but were included in the symptom-based clustering. For the presented numbers, 25 (50 in total) cases for age were missing. For sex, 2 (27 in total) cases were missing. For no. of MDEs, 36 cases were missing. The current SSRI status of 57 subjects in the patient group was not recorded.

### Clinical inventories

All participants completed the BDI-II^20^ and BAI^21^ during recruitment and within 1-2 weeks of the MRI-sessions, comprising 21 items assessing current symptoms. The originally proposed somatic-affective and cognitive factor subscales were used in further analyses. Summary statistics for each item by group are shown in Supplemental Table S1, and a correlation plot with a dendrogram based on hierarchical clustering across all items are shown in Supplemental Figure S1. Largest scores across groups were observed for *lack of energy* (BDI15), *changes in sleeping pattern* (BDI16), *tiredness or fatigue* (BDI20), *nervous* (BAI10) and *indigestion* (BAI18).

### HDDC

BDI-II and BAI symptom scores were z-normalized and submitted to HDDC in the R package *HDclassif*^22^ using standard parameters. HDDC is an unsupervised model-based clustering method based on the Gaussian Mixture Model, and has been shown to outperform similar methods in the R package *mclust*^23^ in terms of accuracy^22^. HDDC also calculates the probability of each subject belonging to each of the clusters, which were used in subsequent analyses. We established the optimal number of clusters using the Bayesian Information Criterion (BIC)^22^, and performed various analyses to assess the robustness and stability of the clustering (see the Supplemental Results) using the *clusteval* R package^24^. To further examine the symptom profiles of each subgroup, based on the full correlation matrix, we assessed the eigenvector centrality of each symptom using the *eigenvector_centrality_und.m* function in the Brain Connectivity Toolbox^25^ in MATLAB R2016B (The MathWorks), yielding a graph-based metric reflecting symptom centrality or importance.

**Table 2.** Demographics of the MRI-subsample

|  | Individuals with<br>no history of<br>depression<br>(n = 72) |  | Individuals with a<br>history of<br>depression<br>(n = 178) |  | *p* |
| --- | --- | --- | --- | --- | --- |
|  | M | SD | M | SD |  |
| Gender (female) | n = 48 |  | n = 127 |  | 0.450 |
| Age | 42.50 | 13.58 | 38.94 | 13.42 | 0.098 |
| Education level (ISCED) | 6.00 | 1.02 | 5.93 | 1.22 | 0.625 |
| Depressive symptoms |  |  |  |  |  |
| Ham-D | 2.89 | 2.11 | 8.03 | 5.78 | <.001 |
| BDI-II | 1.61 | 2.94 | 11.84 | 10.36 | <.001 |
| Anxiety symptoms |  |  |  |  |  |
| BAI | 1.65 | 2.80 | 8.22 | 8.09 | <.001 |
| Other |  |  |  |  |  |
| AUDIT | 4.76 | 3.30 | 6.32 | 5.18 | 0.050 |
| DUDIT | 0.48 | 1.87 | 0.89 | 2.70 | 0.180 |
| Handedness (left) | 7 |  | 6 |  |  |
| History of anxiety disorder | n = 1 |  | n = 53 |  | <.001 |
| History of (hypo)mania | n = 0 |  | n = 30 |  |  |
| No. of depressive episodes | 0 |  | 4.34 | 5.76 | <.001 |
| Currently medicated (SSRI) | n = 0 |  | n = 55 |  | <.001 |
| Currently unmedicated (SSRI) | n = 0 |  | n = 123 |  | <.001 |

2 cases not reported here do not fulfil MDE criteria but have generalized anxiety disorder, but are used in the MRI analysis.

### MRI acquisition protocol

For fMRI analysis a T2* weighted single-shot gradient echo EPI sequence was acquired with the following parameters: repetition time (TR)/echo time (TE)/ flip angle (FA) = 2.500ms/30ms/80°; voxel size, 3.00 × 3.00 × 3.00 mm; 45 transverse slices, 200 volumes; scan time ≈ 8.5 min. Participants were instructed to have their eyes open, and refrain from falling asleep. Scanner noise and subject motion were reduced by using cushions and headphones. For co-registration, we collected a T1-weighted 3D turbo field echo (TFE) scan with SENSE using the following parameters: acceleration factor = 2; TR/TE/FA: 3000 ms/3.61 ms/8°; scan duration: 3 min 16 s, 1 mm isotropic voxels. Due to technical reasons during the time of acquisition, 64 of the individuals were scanned with the initial sagittal phase-encoding (PE) direction and the remaining 186 were scanned with an axial PE direction for the fMRI data.

### Image processing

The FMRI Expert Analysis Tool (FEAT) from the FMRIB Software Library (FSL)^26^ was used for fMRI data processing. This involved brain extraction, motion correction (MCFLIRT),^27^ spatial smoothing (Gaussian kernel, full-width at half-maximum = 6 mm), high pass filtering (100s) and single-session ICA (MELODIC). Estimated mean relative in-scanner head motion (volume-to-volume displacement) was computed with FSL’s MCFLIRT. FMRIB’S ICA-based Xnoiseifier (FIX)^28,29^ was used to automatically classify noise components and regress them out from the main signal, with a threshold of 60. FIX has been shown to substantially improve the temporal signal to noise ratio (tSNR),^30,31^ which was computed before and after FIX.^32^

T1-weighted volumes were skull-stripped using FreeSurfer 5.3^33^ and used for standard space (MNI-152) registration with FLIRT, refining the process with boundary-based registration (BBR)^34^ and FNIRT.

### Group ICA on fMRI data

To avoid bias due to unequal group sizes group-level ICA was performed on a balanced subset of patients and controls (N = 72 in each group).^35^ Model order was fixed at 40, which provides a reasonable trade-off between anatomical sensitivity and specificity.^36^ IC spatial maps and corresponding time-series were estimated using dual regression.^37^ We assessed the spatial maps as well as the frequency profiles following previous recommendations.^38^ We identified and regressed out the time series of 15 noise components, and an additional 6 components (see Supplemental Figure S2) were discarded from further analyses since their spatial maps did not conform with any established resting-state networks or were a mixture between signal and noise, leaving 19 ICs for connectivity analyses.

### Local functional connectivity: sFC and dFC

For sFC, a node-by-node connectivity matrix was created using partial correlations between the time-series, resulting in 171 unique edges. These partial correlations were L1-regularized, with estimated regularization strength (lambda) at the subject level.^35,39,40^

For dFC, the degree of coupling and de-coupling between pairs of brain nodes is conceptualised as the coefficient of variation of delta phi, which is the normalized differences in their wave phases. First, each of the 19 node time-series was narrow-band filtered within 0.04-0.07 Hz, which is required to obtain meaningful phases.^41^ Next, we applied the Hilbert transform, creating an analytic signal, in which we computed the instantaneous phase values for each of the 19 ICs. Lastly, we estimated the Kuramoto order, an index of oscillation between regions at every instant.^42^

### Global-brain level FC

For each individual sFC-connectome we calculated global efficiency, a graph-based measure of topological organization defined as the average inverse shortest path length in a network, using the *efficiency_wei.m* function in the Brain Connectivity Toolbox.^25^ Metastability, a measure of dynamic flexibility whereby the brain transitions through different states, was computed as the standard deviation of the Kuramoto order parameter.^43,44^ Higher metastability is a potential marker for cognitive and behavioural functioning.^43,45,46 47^ Synchrony, a measure of general coherence,^48^ was computed as the mean of the Kuramoto order parameter. It is hypothesized that such coherence allows for the exchange of information within the brain.^49^

### Statistical analyses

Differences between subgroups in between-node (“edge-wise”) sFC and dFC were tested by means of analysis of covariance (ANCOVA) including subgroup, sex, age, PE direction, and mean relative motion. For inference, we used network-based statistics (NBS)^50^ (10000 permutations, α = 0.05), providing control of the family-wise error (FWE) rate on the network-level. Here we tested for main effects of subgroup and the probability of belonging to a specific subgroup on FC. To assess the relative importance of each node, we computed the sum of the test-statistic across all edges. We used a similar approach to test for associations between the BDI-II and BAI sum and factor scores with FC.

We used ANCOVA in R^51^ to independently test for association between subgroup and global efficiency, metastability and synchrony respectively, controlling for mean sex, age, PE direction and mean relative motion. We used the same model to assess the association between the BDI-II and BAI sum and subscale scores with global efficiency, metastability and synchrony independently.

We used Kruskal-Wallis rank sum tests to assess subgroup differences in demographic and clinical variables.

## Results

### Individual clustering using HDDC

HDDC yielded five symptom-based subgroups with differing symptom profiles. Figure 1 shows the mean scores of each symptom for each of the subgroups and the sum scores for the BDI-II and BAI, while Supplemental Figure S3 shows the BDI-II and BAI subscale sum scores. Overall, the subgroups seemed to differ by total severity. However, several other patterns should be noted, especially in terms of symptom centrality (Figure 2). *Unable to relax* (BAI4) was among the most severe symptoms in subgroup 1. *Feelings of dislike* (BDI7), *worthlessness* (BDI14), and *loss in interest* (BDI12) showed highest centrality in subgroup 1, with low centrality for the BAI-symptoms. *Fear of worst happening* (BAI5) was among the most severe in subgroup 2. *Sadness* (BDI1), *feelings of guilt* (BDI5) and *tiredness or fatigue* (BDI20) showed highest centrality in subgroup 2, and the centrality was higher across BAI-symptoms. *Feelings of guilt* (BDI5) was more severe in subgroup 3. *Tiredness or fatigue* (BDI20), *lack of energy* (BDI15) and *loss of pleasure* (BDI4) showed high centrality in subgroup 3. Notably, although the overall symptom severity in subgroup 5 was lower than in subgroup 3, several symptoms were more severe in subgroup 5, and there was an absence of 27 of the total 42 symptoms. Across all subgroups, *change in sleeping pattern* (BDI16) was among the most severe, and was the only symptom present in subgroup 4. Healthy controls and patients were present in all subgroups (Supplemental Figure S4), yet the proportion of patients was higher in subgroups with highest severity scores, specifically subgroups 2 and 1 (X^2^ = 109.69, df = 1, *p* = 2.2×10^−16^). The stability analyses suggest that the clusters were robust, with ≈ 0.75 Jaccard index being the most common for every pair of iterations (Supplemental Figure S5 and S6).

There were no significant differences in sex (X^2^ = 1.42, df = 1, *p* = 0.234) or age (X^2^ = 58.15, df = 53, p = 0.291) between the subgroups in the total sample. For the individuals with a history of depression, there were no significant differences in current SSRI medication status (X^2^ = 0.21, df = 1, *p* = 0.649), comorbidity (X^2^ = 0.32, df = 1, *p* = 0.572), or number of MDEs (X^2^ = 18.72, df = 19, *p* = 0.475) between the subgroups in the total sample. For the MRI subsample, there were no significant differences in sex (X^2^ = 1.04, df = 1, *p* = 0.309), age (X^2^ = 53.41, df = 48, *p* = 0.274), head motion (X^2^ = 252, df = 251, *p* = 0.470), tSNR before FIX (X^2^ = 252, df = 251, *p* = 0.4704), or tSNR after FIX (X^2^ = 252, df = 251, *p* = 0.488) between the subgroups. For the individuals with a history of depression, there were no significant differences in current SSRI medication status (X^2^ = 0.66, df = 1, *p* = 0.416), comorbidity (X^2^ = 0.16, df = 1, *p* = 0.685), or number of MDEs (X^2^ = 17.57, df = 13, *p* = 0. 174) between the subgroups in the MRI subsample.

**Figure 1.**
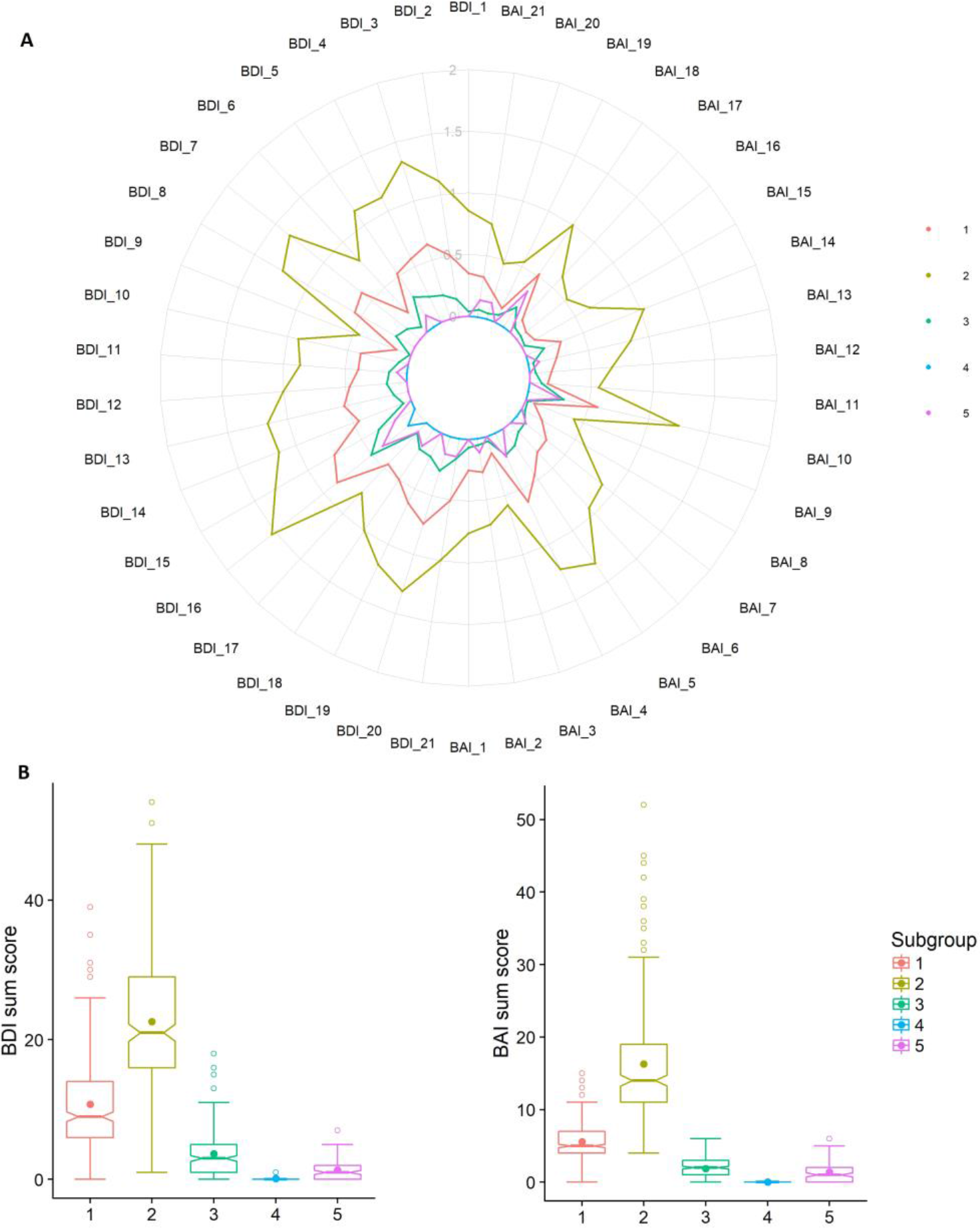
Symptom profiles of the subgroups from HDDC clustering.(A) Mean symptom score of each item of each subgroup. (B) Total BDI (left) and BAI (right) scores for each subgroup.

**Figure 2.**
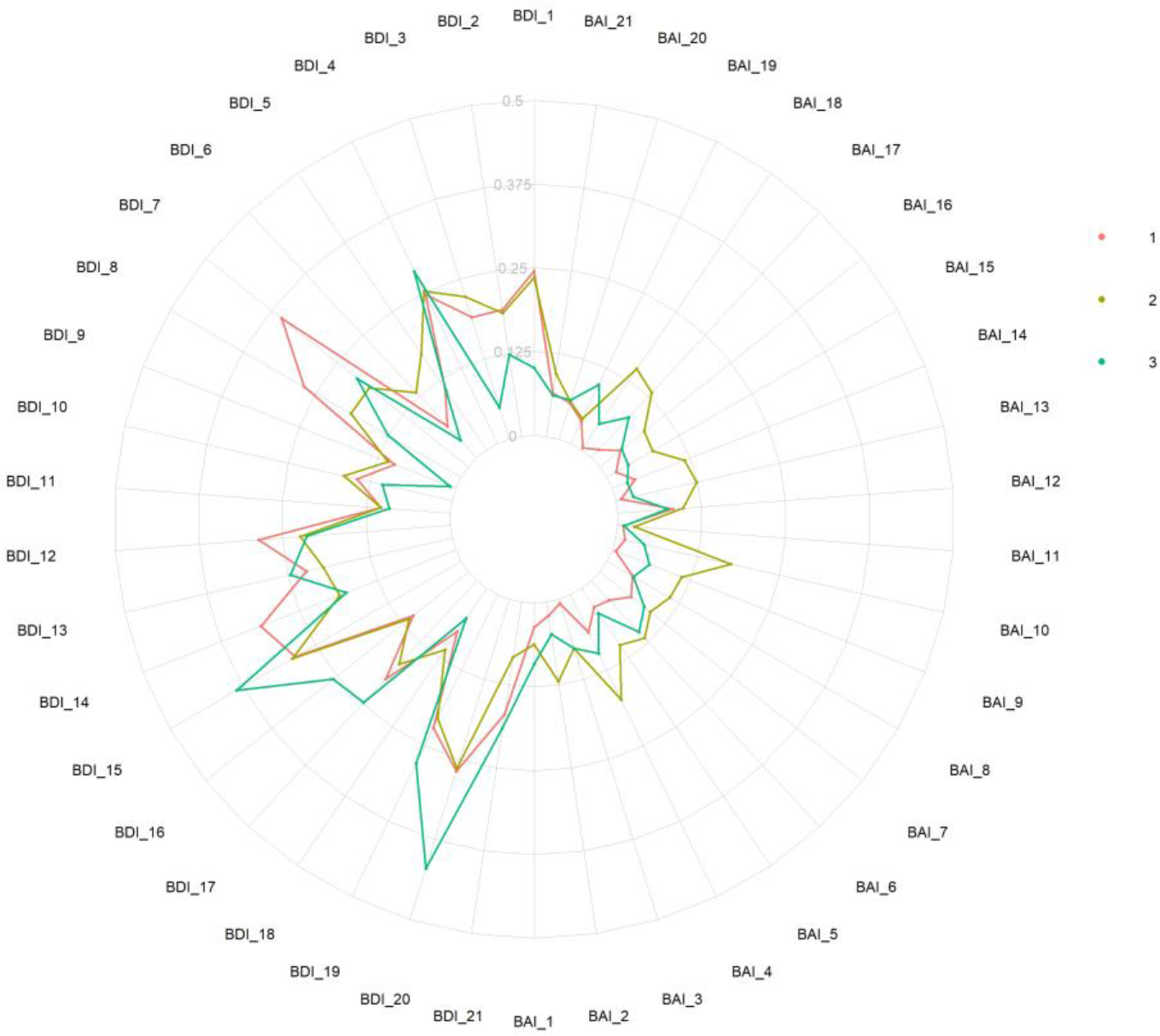
Eigenvector centrality of symptoms for subgroups 1, 2 and 3. Subgroup 4 could not be included because only one symptom, *changes in sleep,* was present. Subgroup 5 was excluded because of the absence of many symptoms (27 of 42) which would change the underlying centrality weighting.

### fMRI-based static FC

NBS revealed a 22-edge subnetwork with significant main effect of subgroup (*p* = 0.033, corrected using permutation testing; Figure 3A). The uncorrected edge level test statistics for this subnetwork are shown in Supplemental Table S2. The strongest differences were seen in edges connecting a default mode network (DMN) component and the fronto-temporal network (IC5-IC16) and between the precuneus and the fronto-temporal network (IC7-IC16; Figure 3B). Figure 3C shows the sum of the test statistics of each node, with largest cumulative effects seen in two default mode network (DMN) components (IC5 and IC6), precuneus (IC7), fronto-temporal network (IC16), cerebellum (IC31) and thalamus (IC39).

NBS revealed a 30-edge subnetwork with significant association with the probability of belonging to subgroup 1 (*p* = 0.015; Figure 3D) and a 24-edge subnetwork with significant association with the probability of belonging to subgroup 3 (*p* = 0.042; Figure 3D). Figure 3E shows the nodes with the largest cumulative effect on the statistical significance of these two subnetworks.

We found no significant associations between BDI-II and BAI sum or subscale scores with sFC (Supplemental Table S3).

**Figure 3.**
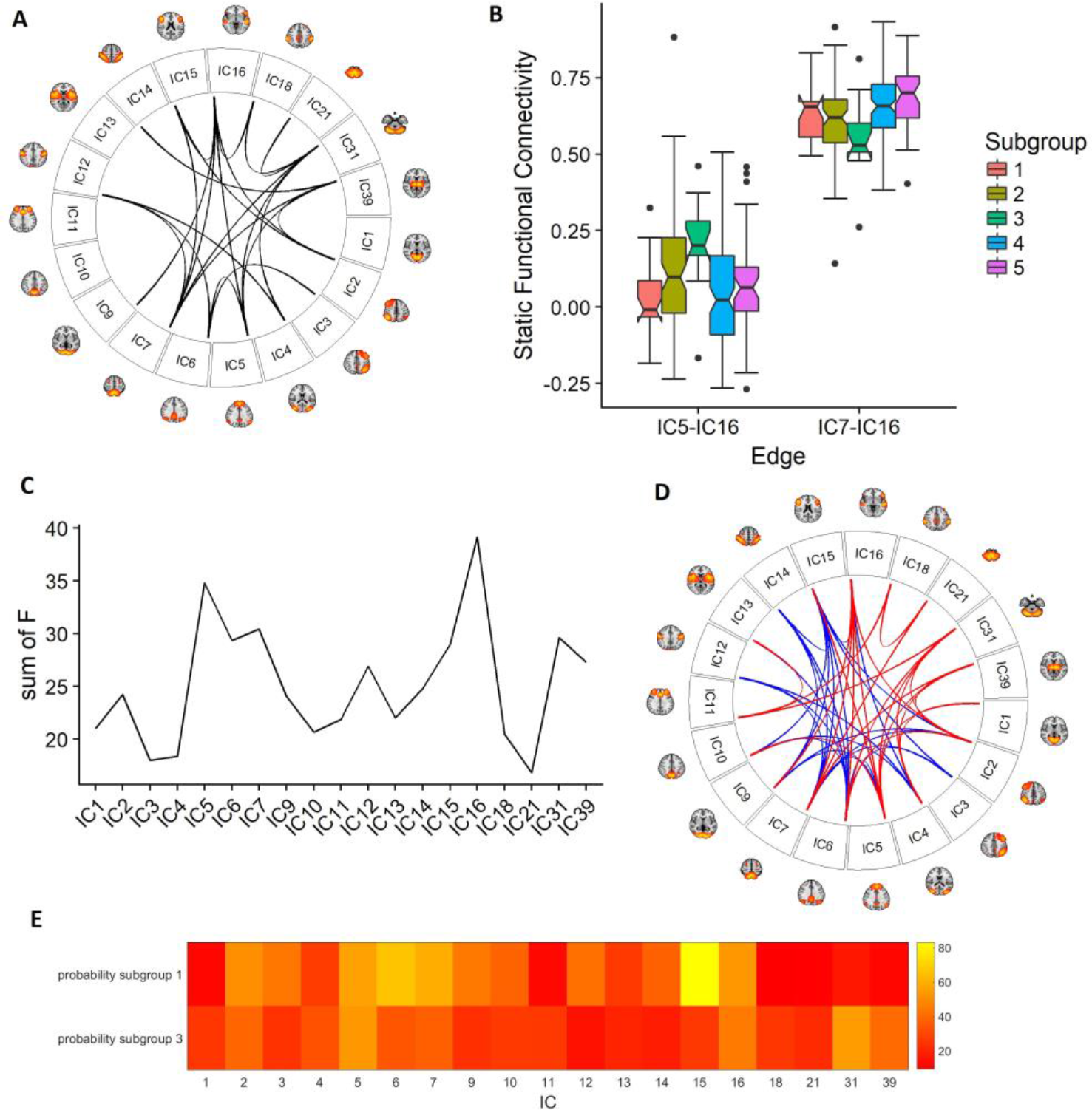
Results from the sFC associations using NBS. (A) subgroup main effect. (B) boxplot of the raw sFC values of the two edges that show the largest main effect of subgroup; between a DMN-component and the fronto-temporal network, and between the precuneus and the fronto-temporal network. Fig. 3C: Sum of test statistic (f-stat) showing the cumulative effect of an IC node on the subgroup main effect. (D) SFC association with the probability of belonging to subgroup 1 (blue) and subgroup 3 (red). (E) Sum of test statistic (f-stat) showing the cumulative effect of an IC node with the association of the probability of belonging to subgroup 1 (upper row) and subgroup 3 (bottom row).

### Dynamic FC

NBS revealed no significant main effect of subgroup on dFC. We found no significant association between BDI-II and BAI sum or subscale scores with dFC (Supplemental Table S4).

### Global-brain level analyses

There was no significant association between subgroup or BDI-II and BAI sum or subscale scores with global efficiency, synchrony or metastability (Supplemental Table S3).

## Discussion

Using high-dimensional clustering of individuals based on current symptoms of depression and anxiety, we have identified five subgroups cutting across diagnostic boundaries in 1083 subjects with a history or no history of depression. Subsequent analysis in a fMRI-subsample revealed a brain sFC pattern with main effect of subgroup, with the fronto-temporal network as a major node. There were no significant associations with conventional symptom domains, supporting that the data-driven clustering provides a more biologically sensitive grouping.

Previous studies have used similar methods to provide data-driven symptom-based stratifications of depression. One study identified a melancholic and a separate atypical subgroup ^52^ which is in line with the DSM-V ^53^. However, this is not a consistent finding across such studies where the most common pattern is total severity differences ^54^ which provides support for a dimensional symptom-based approach. Despite this, the subgroups in the current study exhibit unique symptom profiles both in the pattern of severity and especially in centrality. Notably, subgroup 5 has an absence of many symptoms while the only symptom in subgroup 4 was *change in sleeping pattern,* showing a high degree of specificity. Interestingly, at least one of the three main symptoms that must be present for an MDE in the DSM-V, have different centralities in the subgroups: *sadness* and *loss in interest* have higher centrality in subgroup 1, whereas *loss of pleasure* has higher centrality in subgroup 3. Additionally, subgroup 3 is distinct in that *tiredness or fatigue* and *lack of energy* seem to have much higher centrality than the other subgroups. Incidentally, these two symptoms are grouped together in hierarchical clustering (Supplemental Figure S1).

Data-driven subtyping may have clinical relevance. In a two-year follow up study,^55^ the group with persistent depression had higher centrality in *fatigue or loss of energy* at baseline compared to the remitted group. This symptom specificity could suggest that such subgroups have different underlying mechanisms and environmental triggers. For instance, life stress has been shown to have a substantial impact on interest^56^ whereas romantic breakup was strongly associated with guilt.^57^ *Change in sleeping pattern* is the most severe symptom across all the subgroups, implying that it is more prominent than expected in terms of traditional diagnostic criteria. Recently, different sleep profiles were independently associated with specific patterns of depression comorbidity,^58^ and distinct abnormalities in DMN functioning.^59^

The subgroups showed differential sFC in a range of brain networks, especially involving the fronto-temporal node (IC16). The brain regions encompassing this node are involved with executive functions^60,61^ and external information processing.^62^ The largest edge difference in sFC was between a DMN sub-component and fronto-temporal node (IC5-IC16), which is one of the most consistent FC finding in depression^9^ and can indicate negative self-referential processes.^63,64^ Another edge that exhibited strong sFC differences was between the precuneus and a fronto-temporal node (IC7-IC16). Activity within the precuneus has been associated with increased number of depressive episode^65^ and rumination ^66^. Two other implicated nodes were the cerebellum (IC31) and the thalamus (IC39). Lower cerebellar volume has been associated with decreased emotional memory ^67^, whereas thalamic volume reduction has been associated with deficits in top-down regulation of negative emotions in depression ^68^.Intriguingly, we observed unique sFC patterns associated with the probability of belonging to subgroups 1 and 3, with only a 5-edge overlap. Here, subgroup 3 was uniquely associated with sFC in the supramarginal (IC18), motor (IC21), cerebellar (IC31) and thalamic (IC39) nodes, while subgroup 1 was associated with a higher cumulative effect of the inferior-midfrontal node (IC15).

We found no differences in dFC, global efficiency, metastability or synchrony between the subgroups. We found no significant association between any of the symptom scores with any of the FC measures. Taking these findings together, the sFC associations with the subgroups are partly explained by specificity of symptom profiles beyond total severity differences. Therefore, we argue that a symptom rather than a syndrome-based approach is better suited for elucidating depression symptom heterogeneity.

Two recent studies have identified “biotypes” of depression based on sFC. Drysdale and colleagues^69^ identified four biotypes, whereby biotypes 1 and 2 are similar to subgroup 3 in terms of fatigue, biotype 3 is similar to subgroup 1 in terms of interest, while biotype 4 is similar to subgroup 2 in that anxiety is prominent. The most important features in these biotypes were frontostriatal network dysfunction coupled with anhedonia, and limbic network dysfunction coupled with anxiety. Intriguingly, these subgroups responded differentially to an experimental transcranial magnetic stimulation treatment, showing the potential clinical utility of such subgroups. Price, Gates, Kraynak, Thase and Siegle^70^ identified one biotype characterized by typical DMN connectivity, and a second biotype with increased dorsal anterior cingulate connectivity with higher rates of anxiety and consisted predominantly of females. Both studies and the current study highlight the importance of anxiety in depression, suggesting some convergence across FC and symptom-based clustering. However, FC-based clustering methods are novel, needing validation and replication in independent studies. A strength of the current study is a more detailed range of symptoms.

One limitation of this study is that we included few severely depressed patients, which may have biased the results towards the less severe end of the spectrum. Further, the patient group was clinically heterogenous, with differing history of (hypo)mania, current SSRI medication use and number of depressive episodes. However, there were no significant subgroup differences on any of these factors. Moreover, a recent large-scale meta-analysis of depression studies^6^ found no differences in fMRI results when accounting for such factors.

Methodological variability may account for the discrepancy in previous fMRI findings,^16,71,72^ e.g. related to the definition of the nodes (e.g. ICA vs. ROI-based) and edges (e.g. full vs partial correlations). Based on graph-theoretical accuracy, ICA has been shown to outperform ROI-based node definition, and ROIs may not conform well with functional and anatomical boundaries.^73,74^ Sliding-window analyses are the most common method of analysing dFC, but one issue is unsuitability for fMRI sequences that are less than 10 minutes.^75^ Head motion is a major confounder in FC studies,^76,77^ but this was taken into account in the analyses.

## Conclusion

We identified five robust subgroups with specific clinical symptom profiles. FMRI analysis revealed that these subgroups were characterized by distinct static brain connectivity patterns, in particular implicating a fronto-temporal node. These neurobiologically sensitive subgroups based on a dimensional and symptom-based approach may help move the field towards precision and individualized treatment of depression.

## Acknowledgements

This work was supported by the South-Eastern Norway Regional Health Authority (2014097, 2015073), the Research Council of Norway (249795), and the Department of Psychology, University of Oslo.

We thank Ragnhild Bø and Martin Aker for providing clinical data from their respective research projects. We thank Grethe Løvland and Svein Are Vatnehol for assistance with the technical aspects of the MRI protocol. Finally, we thank Dani Beck for substantial contribution of the MRI data acquisition.

